# Temporal Regulation of Cold Transcriptional Response in Switchgrass

**DOI:** 10.1101/2022.08.01.502344

**Authors:** Thilanka Ranaweera, Brianna N.I. Brown, Peipei Wang, Shin-Han Shiu

## Abstract

Switchgrass low-land ecotypes have significantly higher biomass but lower cold tolerance compared to up-land ecotypes. Understanding the molecular mechanisms underlying cold response, including the ones at transcriptional level, can contribute to improving tolerance of high-yield switchgrass under chilling and freezing environmental conditions. Here, by analyzing an existing switchgrass transcriptome dataset, the temporal *cis*-regulatory basis of switchgrass transcriptional response to cold is dissected computationally. We found that the number of cold-responsive genes and enriched Gene Ontology terms increased as duration of cold treatment increased from 30 min to 24 hours, suggesting an amplified response/cascading effect in cold-responsive gene expression. To identify genomic sequences likely important for regulating cold response, machine learning models predictive of cold response were established using *k*-mer sequences enriched in the genic and flanking regions of cold-responsive genes but not non-responsive genes. These *k*-mers, referred to as putative *cis*-regulatory elements (pCREs) are likely regulatory sequences of cold response in switchgrass. There are in total 655 pCREs where 54 are important in all cold treatment time points. Consistent with this, eight of 35 known cold-responsive CREs were similar to top-ranked pCREs in the models and only these eight were important for predicting temporal cold response. More importantly, most of the top-ranked pCREs were novel sequences in cold regulation. Our findings suggest additional sequence elements important for cold-responsive regulation previously not known that warrant further studies.

## INTRODUCTION

Switchgrass (*Panicum virgatum L*.) is a perennial C4 grass species native to North America and identified as a major lignocellulosic feedstock for biofuel production (Sanderson et al., 2006). Higher biomass production has been a major breeding target and a potent research area in switchgrass. However, high-yielding switchgrass cultivars grow in narrow climatic niches and are known to be less productive under drought, high salinity, and freezing/chilling environmental conditions (Lovell et al., 2021; Sage et al., 2015; Zhuo et al., 2015). Expanding the growing range of high-yielding switchgrass cultivars has been proposed as a way to achieve economic bioenergy production (Sanderson et al., 2006). Coupling high biomass production with low and freezing temperature tolerance can be an effective way of increasing the range expansion of high-yielding switchgrass cultivars. Thus, it is important to understand which genes and how they are responsive to cold stress in cold-resistant switchgrass cultivars.

The ability to tolerate and/or resist cold stress has been an active area of research with respect to the underlying genes, their transcriptional regulators, and signaling pathways (Manasa et al., 2021; Park et al., 2018; Thomashow, 2010). At the level of transcriptional regulation, the C-repeat-binding factor (CBF) cold response pathway is one of the best characterized. In *Arabidopsis thaliana*, three *C-Repeat Binding Factor/Dehydration Responsive Element-Binding Protein 1* (*CBF/DREB1*) transcription factor (TF) genes are rapidly up-regulated in response to cold stress (Liu et al., 1998; Stockinger et al., 1997). Such rapid cold response is due to a signaling network that is active upon cold stress. During cold treatment, cellular Ca^+2^ is elevated and activates Calmodulin proteins (CAMs). CAMs then bind to promoters of *CAM-binding Transcription Activators* (*CAMTAs*) and up-regulate expression of *CAMTAs*. Finally, CAMTAs bind to the conserved CGCG-box in *CBF* genes and up-regulate their transcription. Another well-studied regulator of *CBF* expression is the *Inducer of CBF Expression* (*ICE*) (Chinnusamy et al., 2003). ICE TFs are activated through low temperature mediated sumoylation and subsequently bind to ICE-box promoters in *CBF* genes to activate its transcription (Chinnusamy et al., 2010, 2007, 2003). CBF TFs then up-regulate over 100 cold regulated (COR) and low-temperature induced genes by binding to C-repeat/dehydration-responsive (CRT/DRE) elements, located in promoters of COR genes (Thomashow, 2010). This regulatory hub is known as the CBF regulon which is a major mechanism of cold stress response regulation in plants.

Beyond the CBF regulatory hub, there are examples of other, non-CBF regulatory pathways important for cold stress response in plants. Studies using CBF mutants have shown that TFs rapidly responsive to cold, such as HSFC1, ZAT12, and CZF1, also regulate COR gene expression, indicating CBF-independent regulation (Liu et al., 2019; Park et al., 2018). Another example is BZR1 TFs in the brassinosteroid (BR) signaling pathway that become dephosphorylated upon exposure to cold stress and bind to BR responsive element and E-box in the promoter regions of COR genes such as *WRKY6, SAG21*, and *SOC1* (Li et al., 2017). It is also shown that cold-induced, Abscisic Acid modulated COR gene expression is also shown to work independently from CBF regulon (Liu et al., 1998). There are likely other, non-CBF regulatory mechanisms for plant cold-responsive transcription that remain to be discovered. In addition, in switchgrass, it remains unclear how temporal regulation of cold response is regulated, CBF-dependent or not.

Computational approaches are powerful tools in the identification of genome-wide regulatory patterns in plants under biotic and abiotic stress conditions. In switchgrass, co-expression analysis has been used to establish the potential transcriptional regulatory networks in heat, drought, and biotic stress conditions (Hayford et al., 2022; Pingault et al., 2020). Recently, a comprehensive, transcriptomic study on several panicoid grasses, including switchgrass, revealed that machine learning approaches can be implemented to predict cold stress responses of genes within and between species based on nucleotide frequencies in promoter regions of genes, among other features (Meng et al., 2021). Beyond nucleotide frequencies, a similar approach using longer nucleotide sequences (i.e., *k*-mers) can identify putative *cis*-regulatory elements that are regulatory switches of gene expression under cold stress in switchgrass. Such approaches have been applied to identify the regulatory switches of genes under wounding (Liu et al., 2018; Moore et al., 2022), salinity (Uygun et al., 2017), iron excess response (Kakei et al., 2021), heat, and drought stress conditions (Azodi et al., 2020).

In this study, we aim to apply a similar, machine-learning based approach in switchgrass to assess the involvement of CBF-dependent components of cold response regulation and identify other *cis*-regulatory mechanisms. Using an existing cold stress time course transcriptomes of switchgrass (Meng et al., 2021), we first identified temporally cold-responsive genes. To test the extent to which the temporal cold transcriptional response at different cold treatment duration can be explained using potential *cis*-regulatory sequences, we built machine learning models to predict genes that are up- and down-regulated upon cold treatment in the time course experiment using *k*-mers enriched among up- or down-regulated genes. The *k*-mers that were the most predictive for cold-responsive genes were considered putative *Cis*-Regulatory Elements (pCREs) controlling the temporal transcriptional response. To further reveal the regulatory logic behind the temporal transcriptional response, we examined transcription factors that may bind to pCREs, similarity between pCREs to known CREs, as well as functions of the genes that these pCREs are located on. In addition, to understand if there are common mechanisms underlying the transcriptional response at different time points after cold treatment, we assessed if pCREs identified in one time point were similar to the regulatory elements identified in other time points.

## RESULTS AND DISCUSSION

### Temporal transcriptional response in switchgrass under cold stress

Switchgrass genes responsive to cold stress at different treatment time points (0.5, 1, 3, 6, 16, and 24 hrs) were identified using the transcriptome data from Meng et al, (2021) (**S1 table**). We found that the number of cold-responsive genes, regardless if they were responsive to cold at multiple time points or at a specific time point, increased as the duration of cold treatments (**S1A fig**). This observation is consistent with a cascading effect of transcriptional response over time, similar to responses to other biotic (Ikeuchi et al., 2017; Moore et al., 2022; Ren et al., 2008)and abiotic (Joshi et al., 2016; Ohama et al., 2016) stress conditions. This cascading effect could be because the key regulators are activated sequentially during the cold treatment (Ding et al., 2019a; Lamers et al., 2020). Moreover, as expected, more cold-responsive genes tend to be shared between adjacent time points compared with time points apart from each other (**S1A fig**).

To understand what functions the genes that are responsive to cold stress at different time points tend to have, we conducted Gene Ontology (GO) enrichment analysis (see **Methods**, **S1B and C fig**). GO terms relevant to signaling and activity of transcription factors, such as protein phosphorylation and regulation of transcription, were enriched for genes up-regulated at earlier time points (i.e., 0.5 - 3 hrs, **S1B fig**). These early up-regulated genes may act as initial regulators of genes that are responsive to cold at later time points. Consistent with this, it is known that the accumulation of Ca^+2^ as a result of initial cold sensing activates the expression of calcium-dependent protein kinases (CDPKs), which in turn activate transcription factors that regulate downstream cold stress response (Chinnusamy et al., 2010; Knight and Knight, 2012). Moreover, GO terms such as glucan metabolism and trehalose biosynthesis were also found to be enriched at initial time points. These biological processes are known to be important in the initial cold acclimation in Arabidopsis (Maruyama et al., 2009; Miranda et al., 2007). The GO terms enriched in up-regulated genes at later time points (i.e., 6-24 hrs) may involve biological processes that are required to maintain the functionality of the plant under prolonged cold stress. For example, during prolonged cold stress an increase in plant respiration has been observed (Manasa et al., 2021). As a result of elevated respiration, plants tend to accumulate higher amounts of reactive oxidative species (ROS), followed by the transcription of genes that are responsive to oxidative stress (Wei et al., 2022). This is in line with the enriched GO terms for later cold-responsive genes, such as response to oxidative stress and metal ion transport. Thus, the results from GO enrichment analysis are also indicative of the cascading effect of temporal transcriptional response under cold stress in switchgrass, where initial responsive genes activate later cold-responsive genes that are involved in different physiological and metabolic processes to withstand cold stress conditions.

### Putative *cis*-regulatory elements (pCREs) regulating temporal cold stress responses

The cascading effect of temporal transcriptional response that we observed, as well as the differences between GO terms enriched in genes that were up-regulated at different time points, indicates that the transcriptional regulation differs among time points after cold treatment. To understand how cold-responsive genes are regulated at the *cis*-regulatory level, we first identified *k*-mers in the promoter and gene body regions that were enriched among cold-responsive genes at each time point. Then the enriched *k-mers* were used to establish a predictive model to distinguish cold-responsive genes from non-responsive genes for each time point with machine learning (see **Methods**; **Fig. 1A**). We calculated F-measure (F1 score) on the validation and test instances (held out before model training, see **Methods**). In our modeling setup, the F1 score ranges between one and zero, where one represents a model with perfect prediction, while a score ~0.5 indicates a model with predictions no better than random guesses. Among models distinguishing genes that are significantly up- or down-regulated from non-responsive genes at different time points, the F1s were all higher than random expectation (> 0.7) (**Fig. 1B**), indicating that the sequence information (i.e., *k*-mers) was predictive of cold stress response at a time point.

**Figure 1:**
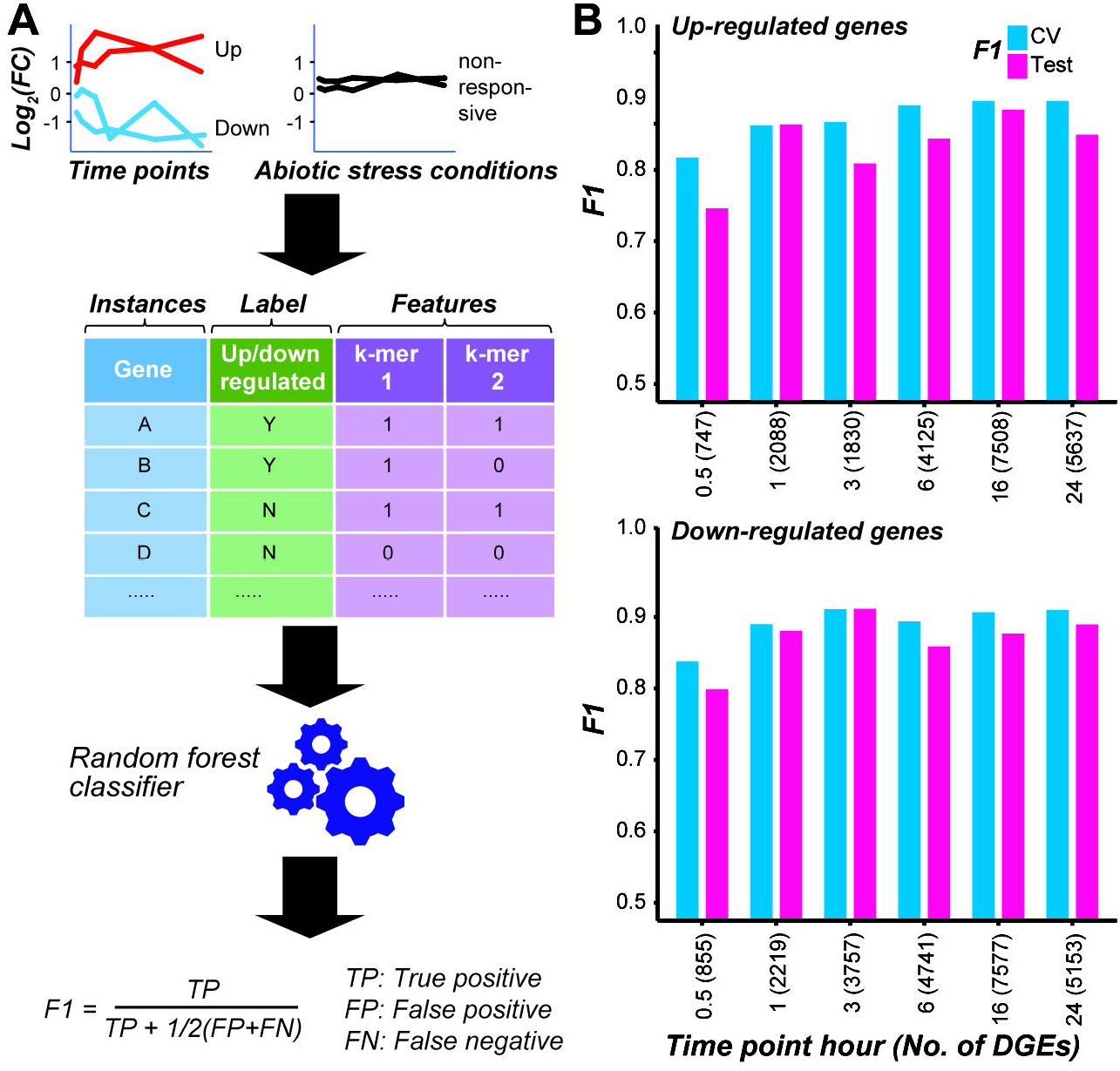
Models predicting the cold responsiveness. (**A**) The overall procedure to model transcriptional response. Genes that are significantly up- or down-regulated at a cold treatment time point were used as positive examples, while genes not responsive to cold treatment at any time point and to other abiotic stresses (dehydration, salt, and drought) were used as negative examples. *k*-mers enriched in the gene body and flanking non-genic regions of the cold-responsive genes were used as predictors (features). RandomForest classifier was used to train models, and the model performance was evaluated using the F1 score. (**B**) Model performances (F1) on the cross-validation (CV) and test sets for each time point model distinguishing genes that were up- (top chart) or down-regulated (bottom chart) after cold treatment for a specific duration from non-responsive genes. The number of positive example genes used in each time-point model is shown in the parenthesis.

Next, we asked what features (*k*-mers) were most predictive of the temporal cold stress response of genes with feature selection. By assessing the model performance improvement by adding features successively from the most to the least important, the minimal number of features required to reach 95% of the optimal model performance was identified for each time point model (**S2 fig**). The *k*-mers that met this criteria for each time point model were defined as pCREs (**S2 and S3 tables**). From here onwards, we focus on the pCREs predictive of up-regulated genes. Some of these pCREs were general across time points (**Fig. 2A**), which may indicate: (1) the genes regulated by these pCREs are responsive to cold across time points; and/or (2) different genes that are responsive to cold stress at different time points are regulated by the same pCRE set. We should note that only 154 and 411 genes for up- and down-regulation across >4 time points, respectively. On the other hand, 16,414 and 16,911 genes are up- and down-regulated in >=1 time points. Considering that very few genes are commonly responsive across multiple time points, the first possibility is unlikely. Some other pCREs were time point-specific (**Fig. 2A**). The remaining pCREs were identified by models predicting genes up-regulated at 2~5, most of the time, disjointed time points (**S3 fig**).

**Figure 2:**
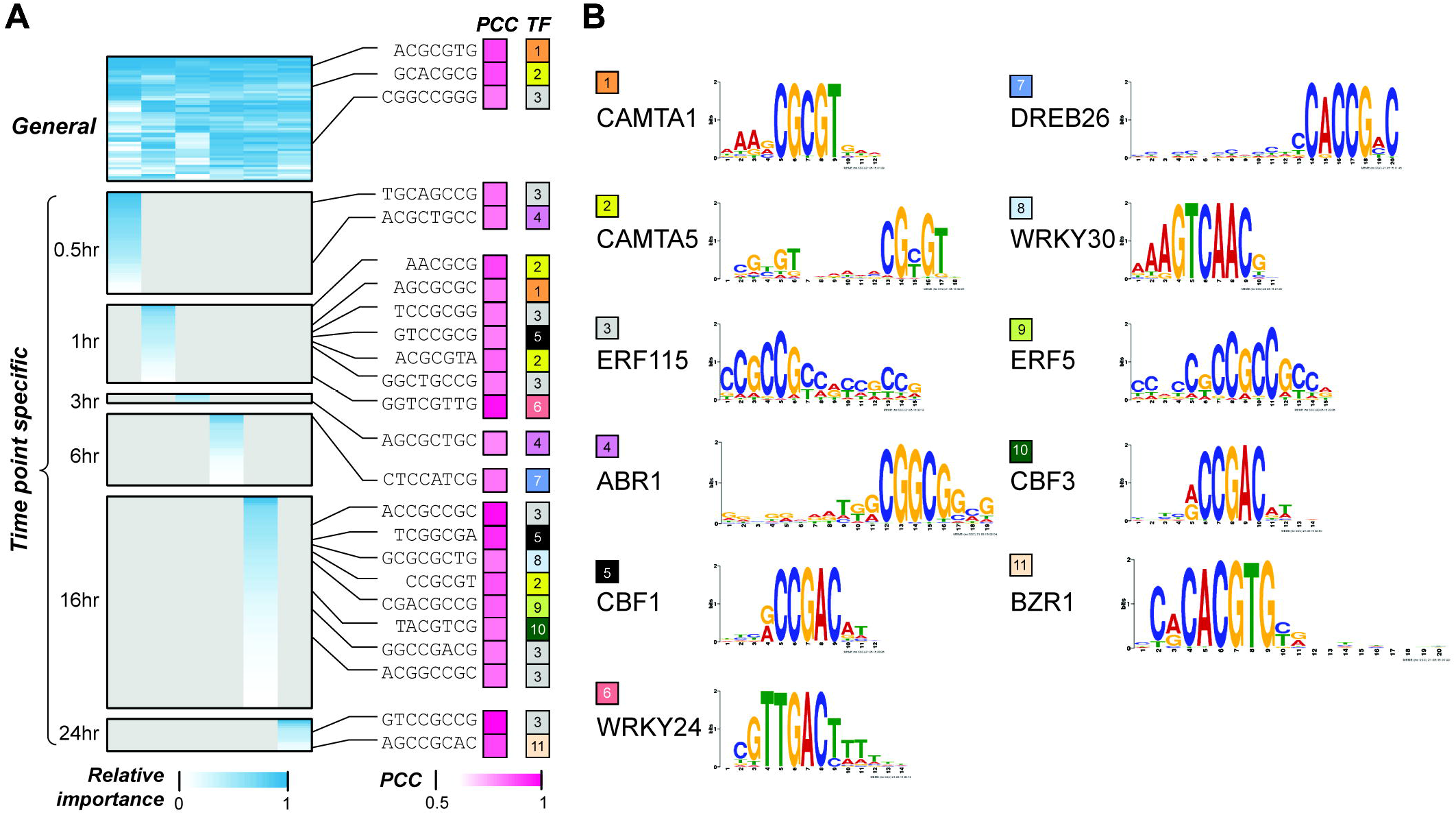
Interpretation of the temporal cold-responsiveness prediction models. (**A**) General and time point-specific pCREs and their similarities to known cold CREs. Heatmap in the left panel shows the relative importance of pCREs, short sequences in the middle indicate pCREs that are similar to CREs known to regulate cold response (cold-TFBMs), which are shown in the right panel. Color scale in blue represents min-max scaled Gini index calculated for features in a time point model; color scale in pink indicates similarities between pCREs and cold-TFBMs. (**B**) Transcription factors (TFs) that bind to the cold-TFBMs are shown with different colors, and the sequence logos of TF binding sites are shown in the rightmost panel. PCC: pearson correlation coefficient. TFBM: transcription factor binding motifs.

### Known cold response regulation transcription factors likely bound to pCRE sites

Previous studies have shown that there are some conserved CREs that control the expression of both early responsive transcription factors (TFs), such as CBF, and downstream cold-responsive genes (e.g., COR genes) that carry out the cold stress tolerance in plants (Chinnusamy et al., 2010; Ding et al., 2019b; Park et al., 2018; Thomashow, 2010). To see if our models have identified binding sites for these known regulators as well as novel CREs, we examined the similarities between the general and time point-specific pCREs and 35 known transcription factor binding motifs (TFBMs) in Arabidopsis using DAP-seq (O’Malley et al., 2016) and CISBP (Weirauch et al., 2014) datasets (**S3 table**). In addition, we collected 35 known TFs regulating plant cold stress response that have binding site information (**S4 table**). Some pCREs that are significantly more similar (see **Methods)** to binding sites of 11 out of 35 known TFs regulating cold response than the 95 percentile of TFBMs from TFs of the same families (**Fig. 2A**, see **Methods**). Two general pCREs were similar to the binding sites of CAMTA1 and CAMTA5 (orange and yellow in **Fig. 2B**). CAMTAs are known to be up-regulated by the activation of Ca^+2^dependent Calmodulin due to cold-induced Ca^+2^ spike (Finkler et al., 2007; Manasa et al., 2021). In addition, CAMTAs are major regulators of *CBF* genes that are known regulators of cold responses, for the immediate cold stress response (Finkler et al., 2007). Consistent with the involvement of CAMTAs in early cold response, pCREs the most closely related to CAMTA binding motifs had the highest feature importance in the 30 min model (CAMTA1 and CAMTA5 ranked 17 and 6, respectively). We should point out that the CAMTA1/5 binding motif-like pCREs were also found in 1hr- and 16 hr-specific sets, indicating that, like in Arabidopsis (Doherty et al., 2009) the CAMTAs may also be involved in maintaining *CBF* or other cold response gene expression that are critical for overall cold acclimation in switchgrass. Because only 11 of 35 cold CREs of known plant cold stress TFs have similar binding sites to general and specific pCREs (**Fig. 2A**), we next examined if they could be recovered using pCREs important in >1 time points (non-specific pCREs, **S3 table**). We found that no new cold CREs can be recovered. Thus, in later discussion, we mainly focus on general and time point-specific pCREs only.

Another notable finding is that pCREs are similar to ERF binding sites (gray and green in **Fig. 2A and B**) and were identified both in the general and most of the time point-specific pCRE sets (excluding the 3 and 6-hrs). Like CBF/DREB TFs, ERF TFs are members of APETALA2/Ethylene Responsive Element Binding Protein (AP2/EREBP) gene family which are known to be involved in multiple stress tolerance (Dey and Corina Vlot, 2015; Park et al., 2021). ERF115 prevents water deprivation in rice under extreme temperatures and drought conditions (Park et al., 2021). Dehydration is a condition that can occur under cold stress and transgenic switchgrass with higher water retention also has an increased cold tolerance (Xie et al., 2019). Despite the lack of experimental evidence for the function of ERF TFs in switchgrass, our findings suggest that ERF TFs may play important roles in cold tolerance in switchgrass. Moreover, there were also pCREs that are similar to binding sites of TFs from other TFs families, such as WRKY, BZR and ABR. pCREs similar to binding sites for BZR1 (rank 1 to 4), WRKY24 (rank seven to eight), and WRKY 30 (rank seven) were also among the most predictive cold-CREs in cold-TFBM models (**S4 fig**). These TFs are known for cold signal transduction and cold stress tolerance via CBF-independent pathways (Park et al., 2015; Ramirez and Poppenberger, 2020). BZR1 is known to be involved in cold stress tolerance through processes such as ROS scavenging (Ramirez and Poppenberger, 2020) and facilitating structural changes in cell membranes and cell walls (Benatti et al., 2012). Moreover, WRKY TFs are also known to be involved in phytohormonal-induced signal transduction for low-temperature tolerance in plants (Park et al., 2018, 2015). ABR1 on the other hand is known to regulate stress responses including cold stress in a CBF-independent, CBL9-CIPK3-mediated, ABA-signaling cascade (Pandey et al., 2005). These findings indicate that our prediction models can not only predict cold-responsiveness for different time points after cold treatment, but also recover known plant cold-TFBMs.

### Potentially novel cold *cis*-regulatory sequences in switchgrass

While known TFs involved in cold-responsive regulation can be identified, 45 pCREs either resembled known TFBMs but the TFs were not known to be involved in cold-regulation. Perhaps more importantly, another 598 pCREs did not have significant similarity to known TFBMs. This raises the question if these pCREs not resembling cold-TFBMs, represent novel component of switchgrass *cis*-regulation under cold treatment. To address this, we compared the informativeness of pCREs identified by our models and the experimentally validated cold-TFBMs for predicting cold stress response. Based on literature search, 35 TFs involved in cold response regulation with binding site information in different plant species (**S4 table**) were used to build models (hereafter referred to as cold-TFBM models). We found that the cold-TFBM models had far worse prediction performance (median F1=0.66) than models built using all pCREs (median F1=0.85, **Fig. 3A**). Since these 11 of 35 cold-TFBMs are significantly similar to top-ranked pCREs (similarity >95% of randomly expected matches, see **Methods**), it is not particularly surprising that the cold-TFBMs predictive of cold responsiveness at different specific time points are similar to the findings in **Fig. 2**, By looking at the feature importance of the cold-TFBMs models built for each of the time points (**S4 fig**), TFBMs of CAMTA1/5 and CBFs were among the most predictive features among the cold-TFBMs time point models.

**Figure 3:**
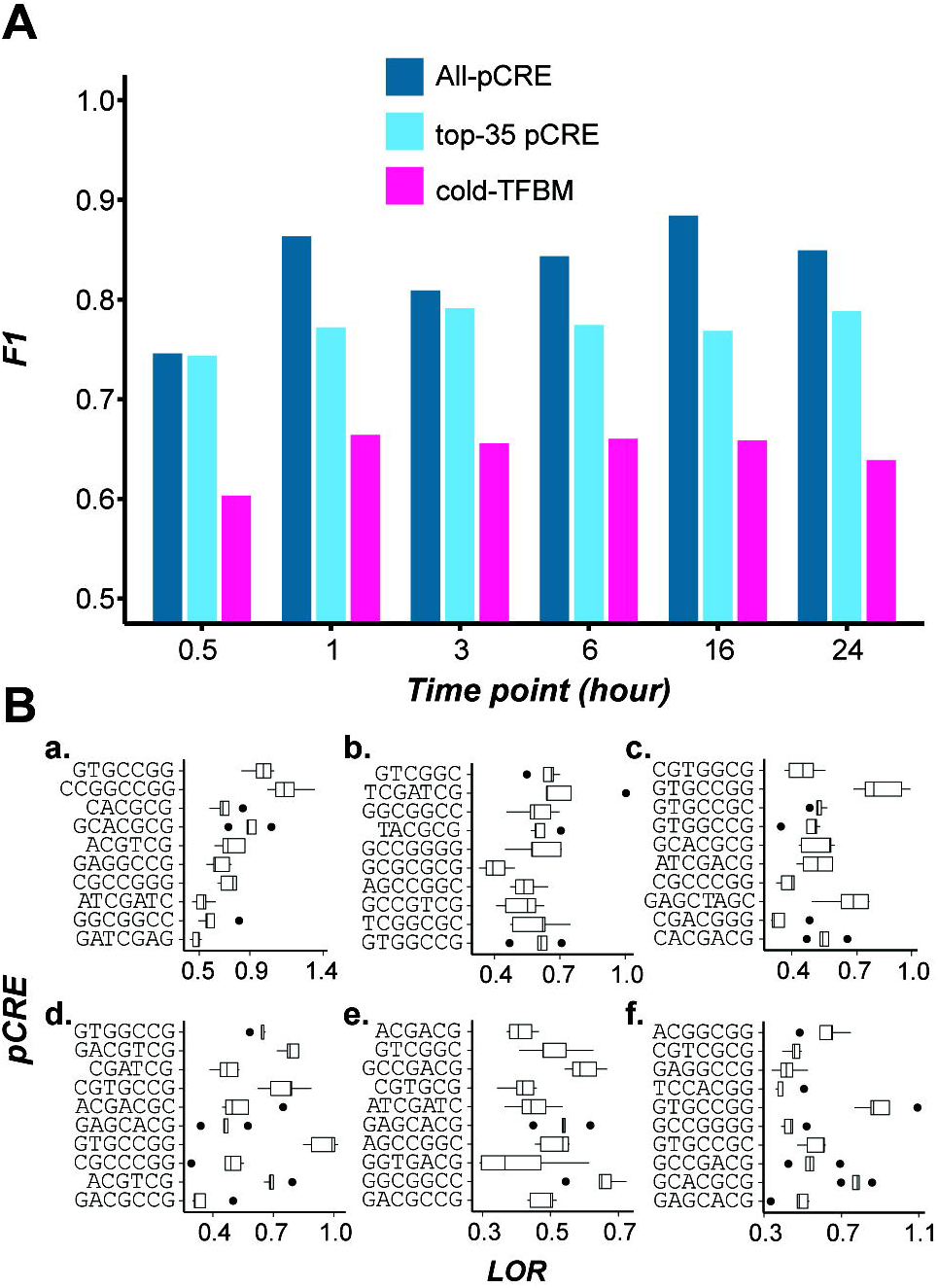
**(A)** Model performance comparison among models built using all the pCREs (blue), top 35 most important pCREs (cyan), and 35 known cold-TFBMs (hot pink). (**B**) Enrichments of top 10 pCREs in 0.5, 1, 3, 6, 16, 24 hr time point models (a-f respectively).

While the all-pCRE models overall performed significantly better than cold-TFBM-based ones (T-test, *p* < 0.01, **Fig. 3A**), it is possible that the all-pCRE models simply have far more features. To address this, we also built models using the top 35 most important pCREs (based on the feature importance of time point models) for comparison. We found that the cold-TFBM models remain worse than models built using the top 35 pCREs (median F1=0.77, *p*<0.01, **Fig. 3A**). This finding, together with that based on all-pCRE models, suggests that pCREs identified in our models contain potentially novel cold-responsive CREs that may or may not be specific to switchgrass. In **Fig. 3B**, the top 10 ranked pCREs from each of the time point models are shown with emphasis on novel pCREs. These novel pCREs are significantly enriched (multiple testing corrected, *p*<0.05) in cold stress up-regulated genes at each time point (Median log odds ratio=0.55). Taken together, the comparison between cold-TFBM models and the all-pCRE or the top-35 pCRE models shows that known cold-TFBMs could not explain cold responsiveness at any particular time point as well. These findings suggest that there are novel temporal *cis*-regulatory components of cold transcriptional response.

### Relationships between pCREs across time points

The majority of top pCREs are sequences that do not resemble TFBMs associated with cold regulation. To further understand how these pCREs we identified may be involved in temporal cold stress regulation, we examined: (1) the similarity of the pCREs across time point models (**Fig. 4A**); (2) importance of pCREs from different clusters in predicting cold response (**Fig. 4B**); (3) functions carried out by the genes that the pCREs were located (**Fig. 4C**); (4) sequence similarities between pCREs and TFBMs (earlier the focus was only on cold-related TFs, **Fig. 4D**); and (5) expression profiles of genes that the pCREs were located (**Fig. 4E**). First, we categorized the pCREs into clusters by calculating the pairwise PCC distance (1-PCC) based on their sequences (see **Methods**; **S5 fig**). The clusters were defined using the same PCC distance threshold as in (Liu et al., 2018), where pCREs with PCC distance <0.39 were considered to be bound by TFs of the same family. The pCREs were grouped into 27 clusters and pCREs in 25 clusters were shared by >1 cold treatment time points. Since pCREs in a cluster are likely bound by TFs of the same family, this finding indicates the involvement of most TF families across time points. These clusters consisted of pCREs important in >1 time points were referred to as non-specific pCRE clusters (**Fig. 4A**).

**Figure 4:**
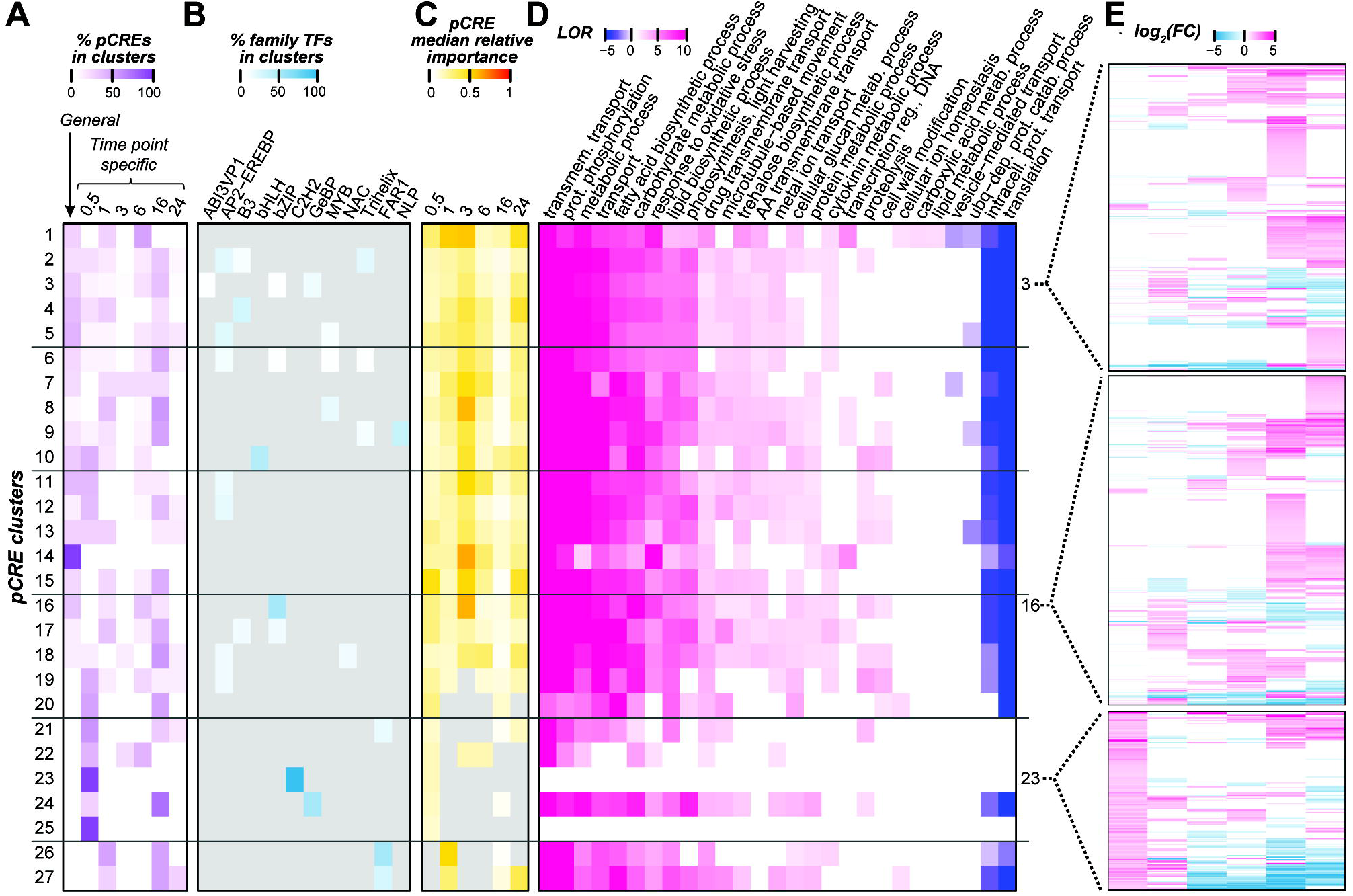
Properties of pCRE clusters which were defined based on sequence similarity. (**A**) Heatmap showing the distribution of general and time point-specific pCREs within a cluster. Color scale represents the percentage of general and time point-specific pCREs in each pCRE cluster. (**B**) Median importance of pCREs in a cluster. Cell color depicts median min-max scaled Gini index of the pCREs within each cluster. Gray color indicates that the pCRE is not used in the time point model in question. (**C**) Potential transcription factors (TFs) that could bind to pCREs in pCRE clusters based on the similarity between pCREs and TF binding sites (TFBS) information based on in-vitro binding assays. A TF was considered to bind a pCRE only if the PCC similarity between the pCRE and its binding sites was above the 95^th^ percentile of the background PCC distribution, which was calculated among TFs in the same TF family. TF families that don’t fall under this threshold were marked in gray. Color scale represents the percentage of pCREs within a pCRE cluster that showed significant similarity with TFBS. (**D**) Significantly enriched biological GO terms of genes containing pCREs in a pCRE cluster. Color scale represents the log_10_(odds ratio), for details, see **Methods**. (**E**) Differential expression of genes that contain pCREs in clusters 3, 16, or 23 at different time points. Each row shows the profile of a gene, and color scale indicates log_2_(FC).

To assess if pCREs in different clusters may regulate distinct sets of genes, we compared the differential expression profiles of genes that contain pCREs from different clusters in different time points (**Fig. 4E** and **S6 fig**). To facilitate interpretation of the differential expression profiles, we encoded the transcriptional responsiveness of a gene at a time point as U, D, N if it is significantly up-regulated, significantly down-regulated, and not differentially expressed, respectively. For example, a profile of “UUDDNN” indicates that the gene is significantly up-regulated at 30 minutes and 1 hr, down-regulated at 3 hrs and 6 hrs, and not differentially expressed at 16 hrs and 24 hrs after cold treatment. Using this strategy, we investigated the frequency of differential expression profiles of genes with pCREs in different pCRE clusters. NNNUUN, NNNUUU, and NNUUUU were the top three most frequent expression profiles found on the genes that contain pCREs in all 25 non-specific pCRE clusters **(S7 fig)**. Because the up-regulatory patterns were contiguous after 3hrs of cold treatment, regulatory switches common between time points may have a role in the up-regulation and maintaining the expression of genes at later time points. Similarly, previous studies also show that in both CBF-dependent and independent pathways, immediately cold-responsive TFs are responsible for up-regulating and maintaining the expression of a large number of downstream cold-responsive genes by binding to conserved regulatory sequences (Li et al., 2017; Park et al., 2015; Thomashow, 2010). Some genes harboring pCREs from non-specific pCRE clusters also had unique expression profiles (expressed in a single time point) as well as much more complex expression profiles (up- or down-regulated in multiple, non-contiguous time points) (**S6 and S7 fig**).

In addition to non-specific clusters, there were two 30 min-specific pCRE clusters (clusters 23 and 25) (**Fig. 4A**). pCREs in these clusters may regulate initial cold transcriptional response. However, these clusters were significantly enriched (q≤0.05) with the genes that are up-regulated only at the 30-min time point compared to genes that contain pCREs in other clusters (**S8 fig**), For example, in cluster 23, UNNNNN, UNUUNN, UNUNNU, UUUUUU, UUDDDD, and UUNNDD are among profiles with the highest degrees of enrichment. There are ~360 different gene expression profiles that contain pCREs in all 25 of the shared pCRE clusters (**S7 fig**). Thus, the temporal regulation of cold transcriptional response is likely mediated through a combination of general CREs that are important for the entire duration, specific CREs that regulates response at particular time, as well as non-specific CREs that regulate a certain duration (contiguous time points) or complicated expression profiles (e.g., UNUNNU). To assess the functions of genes that contained pCREs from pCREs clusters, we examined which GO terms were enriched with genes containing pCREs in a cluster (**Fig. 4C**). Except for the general enriched GO terms (e.g., metabolic processes), genes containing pCREs of non-specific pCRE clusters were enriched with biosynthetic processes that are involved in cold stress responses (e.g., fatty acid biosynthetic process, lipid biosynthetic process, and trehalose biosynthetic process) and specific metabolic processes (e.g., response to oxidative stress, carbohydrate metabolic process) (**Fig. 4C**). These GO terms are known to be enriched in late responsive genes under cold stress (Manasa et al., 2021). Our findings suggest that some genes containing pCREs from these non-specific pCRE clusters may contribute to metabolic processes crucial for cold tolerance. None of the GO terms were enriched for genes containing pCREs in the specific pCRE clusters 23 and 25, potentially due to the small sample size of these two clusters.

### Cold stress regulatory pCREs that do not resemble known TFBMs

To further assess the regulatory role of the pCREs in pCRE clusters, we asked what TFs may bind to these pCREs using the in-vitro TFBM information of 344 Arabidopsis TFs. Although the Arabidopsis and the switchgrass lineages diverged ~200 million years ago (Wolfe et al., 1989), the TFBMs of dicot and monocot TFs from the same families are highly similar (Weirauch et al., 2014). A TF was considered to have the potential to bind to a pCRE if the similarity between its TFBM and the pCRE in question was above the 95^th^ percentile of the similarity distribution calculated among TFBMs in the same TF family (see **Methods**). In addition to members of the AP2-EREBP family discussed previously (**Fig. 2** and **4C**), TFBMs of B3, bZIP, MYB, Trihelix, and FAR1 TF families were also found to have a significant similarity to pCREs in multiple clusters (**Fig. 4C**). In soybean, the bZIP TFs are known to regulate cold stress in ABA-dependent pathways by inducing the expression of downstream COR and ERF type genes that help plants to resist cold stress conditions (Liao et al., 2008; Yu et al., 2020). Moreover, in tomatoes, the Trihelix type TFs are known to be up-regulated under cold stress conditions, and activate downstream genes with products that modulate stomatal conductance to prevent water loss (Liu et al., 2012; Yu et al., 2018). In apples, R2R3-MYB TFs were found to be induced by cold stress and activate ROS scavenging genes (An et al., 2018).

Aside from 19 clusters containing pCREs resembling known Arabidopsis TFBMs, eight clusters did not contain pCREs resembling TFBMs we investigated (**Fig. 4B**). These pCREs are referred to as “unknown” pCREs (those with “between” threshold in **S3 table**). In our time-point models, those unknown pCREs were also important for predicting cold responsiveness of a gene (**S3 table**) as indicated by the median importance of pCREs in clusters (**Fig. 4C**). Furthermore, the feature importance ranks of these pCREs in predicting cold transcriptional response in the time point models (median rank=0.45) are significantly similar (T-test, *p*-value<0.01) to those of pCREs resembling known TFBMs (median rank=0.38). Using general pCREs as examples, we built models to predict genes up-regulated at different time points using solely pCREs similar to known TFBMs (n=16), and another model with unknown pCREs (n=38). We found that the performances of models built using general pCREs similar to known TFBM (median F1=0.66) and general “unknown” pCREs (median F1=0.70) were not significantly different (T-test, *p*-value>0.01). This result also suggests that “unknown” pCREs have similar importance to pCREs that resemble known TFBMs in predicting temporal cold-stress response in switchgrass. The reasons we did not find similar TFBMs to these pCREs may be because the threshold we used to assign a pCRE to TFBSs was too stringent. However, the threshold used was established as the degree of similarity that allows binding motifs of a plant TF family to be identified (Azodi et al., 2020). Thus, it was not asking if a pCRE resembled a specific TFBM, but the binding motifs at the level of family. The second reason may be that Arabidopsis TFBMs were used, which may miss TFBMs specific in other species. Although there is broad conservation of TFBMs across species, even between plants and humans (Weirauch et al., 2014), this can only be assessed with additional experimental studies either through DAP-seq or one-hybrid assay. Another possibility is that the Arabidopsis TFBM data may miss binding sites due to the limitations of in vitro binding assays (Bartlett et al., 2017). Finally, it is also possible that, instead of TFBMs, a subset of pCREs may represent motifs relevant for levels of regulation beyond transcription, such as post-transcriptional or translational regulation. This possibility remains to be investigated.

## CONCLUSION

In this study, we aimed to find DNA regulatory switches responsible for temporal transcriptional response in switchgrass under cold stress conditions. By examining the number of cold-responsive genes at different time points, and the functions these genes tend to have, we found a cascading effect of gene transcriptional responses with regards to the time the plant was exposed to cold stress. The *k*-mers enriched for cold-responsive genes at a particular time point were predictive of the cold responsiveness of genes at that time point. By examining the top most predictive *k*-mers, we were able to identify well known CREs that regulate cold stress response in plants, indicating the usefulness of our models. Based on similarity of a subset of pCREs to known cold TFs, switchgrass cold stress response is mediated through both CBF-dependent and independent pathways. Beyond the known cold-responsive CREs, additional pCREs not known to be regulating cold response were identified. Some pCREs were identified in specific time point models, while others (general and non-specific pCREs) appeared to be relevant to regulation of cold response at multiple, sometimes disjoint, time points. In the latter case, differential expression profiles of genes containing these pCREs show complex patterns throughout the time course.

A substantial fraction of the pCREs do not resemble known binding motifs of known cold response regulatory TFs or, in general, Arabidopsis TFs with in vivo binding data. However, the regulatory function of these pCREs in cold responses needs to be experimentally validated using knockout lines and additional efforts, including modeling complex expression patterns under cold stress response (i.e., non-contiguous, up-/down-regulation) to identify the pCREs responsible for complex temporal expression and modeling cold stress response using combinations of pCREs to identify complex expression patterns under cold stress are required to fully understand the cold-responsive *cis*-regulatory code in switchgrass. We also emphasize how building computational methods and their interpretations are important for identifying the global patterns of gene expression and their context-specific regulatory elements. This study provides sequence elements that regulate temporal cold stress response, allows a systematic understanding of the temporal cold stress regulation in switchgrass and, with subsequent validation studies, the information can be used as the bases for fine tuning switchgrass tolerance to cold stress.

## MATERIALS AND METHODS

### Transcriptome data collection, preprocessing, and gene-set enrichment analysis

The switchgrass cold response RNA-seq data were from a published study of a time course (0.5, 1, 3, 6, 16, and 24 hrs) under cold treatment (6 □) with paired control samples (29 □/23 □ in a 12-h/12-h day/night cycle) (Meng et al., 2021). Switchgrass transcriptomes under three other stress conditions were from three published studies [Dehydration ((Zhang et al., 2018)), salt ((Zhang et al., 2021)), and drought ((Zuo et al., 2018)]. The RNA-sequencing (RNA-seq) data of these studies were downloaded from NCBI-SRA database (https://www.ncbi.nlm.nih.gov/sra), processed, and used to generate raw counts and transcript abundance (transcripts per million, TPM) using an RNA-seq analysis pipeline (https://github.com/ShiuLab/RNA-seq_data_processing.git). For mapping RNA-seq reads, *Panicum virgatum* v5.1 genome and the corresponding genome annotations were downloaded from the Joint Genome Institute (JGI) database (https://jgi.doe.gov). Only reads that were uniquely mapped to the genome were used. Differential expression of genes (fold change, FC) contrasting cold stress treatment and corresponding control at each time point and false discovery rate corrected *p*-values were calculated using the EdgeR package implemented in R (Robinson et al., 2010).

Gene Ontology (GO) annotations of switchgrass genes were downloaded from JGI Data Portal as of 07.08.2021 (https://data.jgi.doe.gov). Fisher’s exact test was conducted to identify GO categories enriched in cold-responsive genes at each time point versus all the other genes in the genome. The resulting *p*-values were adjusted using the Benjamini-Hochberg method (Benjamini and Hochberg, 1995), and GO terms with adjusted *p*-values ≤ 0.05 were considered as enriched for cold-responsive genes (https://github.com/ShiuLab/Manuscript_Code/tree/master/2022_switchgrass_cold_pCREs). The GO enrichment analysis was also conducted for genes that contain pCREs from the same pCRE distance cluster versus all the genes in the genome (see next sections).

### Identification of cold-responsive putative *cis*-regulatory elements (pCREs)

Cold-responsive genes were defined as genes that were either significantly up-regulated (Log_2_FC≥1 and adjusted *p*≤0.05) or down-regulated (Log_2_FC≤-1 and adjusted *p*≤0.05) upon cold treatment at each time point. Genes were defined as non-responsive to cold at any of the six time points and nonresponsive to the other three stress conditions mentioned above (|log_2_FC|<0.5 and/or adjusted *p*>0.05). Here, stress conditions other than cold treatment were considered to define non-responsive genes, because previous studies have found that stress-responsive CREs could activate genes under multiple stress conditions (Azodi et al., 2020; Zou et al., 2011). Thus, contrasting the cold-responsive genes against genes that are not responsive to combined stresses would allow us to identify the full scale of pCREs, i.e., both cold-stress-specific pCREs and pCREs responsible for multiple stress conditions including cold stress.

To identify pCREs, we applied a combination of a *k*-mer enrichment approach and machine learning. To avoid data leakage, for each time point, cold-responsive genes (up- or down-regulated after cold treatment) and non-responsive genes were split where 80% of the genes were used as the training set and 20% were the test set. The test set was set aside and was not used for any pCRE identification or modeling steps. For the *k*-mer enrichment step, genes in the training set were further split into five bins. For each bin, we first identified all possible *k*-mers (*k*=5-8 nucleotides where a forward *k*-mer was considered as the same as its reverse complements) from 1kb upstream, gene body including 5’ and 3’ untranslated regions, and 1kb downstream regions of both cold-responsive and non-responsive genes. *K*-mers enriched for cold-responsive genes (Fisher’s exact test adjusted *p*-value<0.05) were identified for each bin, and the *k*-mers commonly enriched among all five bins were used as features to establish machine learning models classifying cold-responsive genes (positive examples) and non-responsive genes (negative examples) in the training set.

To create a balanced training dataset (same numbers of positive and negative examples), genes in the minority class with fewer instances were randomly up-sampled using the Synthetic Minority Over-sampling Technique (Chawla et al., 2002). We also experimented with down-sampling where the majority class was randomly selected to match the number of minority class genes. Classification models were built for each time point to predict cold-responsive and non-cold responsive genes using the random forest algorithm (Breiman, 2001)grid search was conducted based on 60 hyperparameter combinations (‘max_depth’: [3, 5, 10], ‘max_features’: [0.1, 0.5, ‘sqrt’, ‘log2’, None], ‘n_estimators’: [10, 100, 500, 1000]) in a five-fold cross-validation scheme where every gene was used in the validation set exactly once. The optimal hyperparameter set was selected based on F1 score of the validation set predictions. F1 measure is the harmonic mean of precision and recall. An “optimal” model for each time point was then built using all training instances with the optimal hyperparameters. The final model for each time point was then applied to predict the cold responsiveness of genes in the testing set and model performance was evaluated using F1 measure.

### Selection of minimal pCRE sets as features and determining relationships between pCREs

To identify the minimal number of features (enriched *k*-mers) that have a similar performance as the optimal model using all features to distinguish cold-responsive from non-responsive genes, features were selected based on Gini importance defined as the impurity difference of a node in the decision tree when the feature in question is used, a measure of the contribution of a feature for distinguishing the cold-responsive and non-responsive genes. New models use the training set again by increasing the numbers of features used, starting with just the top 10 important features and, for subsequent models, increasing the number of features by 20 in order of decreasing feature importance. The trend line of the cross-validation F1 score against the number of features was fit with the Michaelis-Menten Equation. For each time point the minimal number of features was determined as where the fitted line had a near zero differential (e.g., the 30 min model, **S1 fig**), or where the F1 first reached 90% of the optimal model F1 if there was no clear plateau (e.g., the 30 min model, **S1 fig)**. Features within the minimal set were designated as pCREs for the cold response at the time point in question. To determine the similarity between pCREs, pairwise PCC distances between pCREs were calculated using the TAMO package (Gordon et al., 2005), implemented in R. The distance matrix was used to construct a UPGMA tree using average linkage in the library ‘cluster’ in R (Maechler et al., 2012). Sequence similarity of 0.39 was used as a threshold, such that pCREs with similarity >0.39 can be treated as a single pCRE (Liu et al., 2018). For each cluster of pCREs, the proportion of pCREs in different categories (general or time point-specific pCRE groups) were calculated using custom scripts (https://github.com/ShiuLab/Manuscript_Code/tree/master/2022_switchgrass_cold_pCREs).

### Identification of transcription factors (TFs) with binding sites similar to pCREs

The assessment of sequence similarity between pCREs and known transcription factor binding sites (TFBSs) was carried out using the Motif Discovery Pipeline (https://github.com/ShiuLab/MotifDiscovery.git) as described in (Azodi et al., 2020). For this analysis, only the pCREs responsible for up-regulation upon cold treatment were considered. Known TFBS data was retrieved from two datasets: (1) DNA Affinity Purification sequencing (DAP-seq) database, where *in-vitro* DNA binding assays were performed for 344 TFs in *Arabidopsis thaliana;* (2) Catalog of Inferred Sequence Binding Preferences (CIS-BP) database, where position frequency matrices (PFMs) for TFBS of 190 TFs (non-redundant TFBS with DAP-seq database) in *A. thaliana* were available (Weirauch et al., 2014). To assess the similarity between pCREs and TFBSs, the Pearson’s Correlation Coefficients (PCC) between the position weighted matrices (PWMs) of pCREs and PWMs of TFBSs were calculated as described in (Azodi et al., 2020). The top matching TFBS for each pCRE was reported in three threshold levels (same TF, same family, or significantly more similar than randomly expected) as described in (Azodi et al., 2020). To determine the similarity between pCRE and TFBMs for TFs regulating cold response, we checked if pCRE-TFBM PCC is higher than 95^th^ percentile of the PCCs calculated among TFBMs of different transcription factors families. This is a mid-stringency threshold out of the three thresholds we used to find similarities between pCREs and TFBMs. Since we are using Arabidopsis TFBMs to identify similar binding sites of specific TFs switchgrass we wanted to use TFBMs with the highest similarity when compared with other families of TFs, which with a higher stringency threshold would not have been found. Using this mid-stringency threshold we will be able to say if a pCRE resembles a specific binding site of a particular TF in comparison with other TFs in different TF families.

To assess how well the binding sites of TFs known to regulate cold response might predict cold response, we collected known cold regulation TFs through a literature search (**S4 table**). Using PWMs of binding sites of TFs known to regulate cold stress in plants (cold-CREs), we mapped similar binding sites in up-regulated genes in different time points. Based on absence/presence of cold-CREs in a gene we recreate feature tables for genes that are up-regulated in each time point. Using the similar machine learning methods used in the “**Identification of cold-responsive putative *cis*-regulatory elements**” section, we made models to predict cold responsiveness of a gene up-regulated in each time point using cold-TFBMs. The performance of these models were then compared to our original time point models.

## SUPPLEMENTAL FIGURE LEGENDS

**S1 figure:** Properties of cold-responsive genes at different time points. (**A)** Matrix showing the number of up-regulated (top left triangle) and down-regulated (bottom right triangle) genes at different time points after cold treatment. Color scale and number within the cell on the diagonal represent the count of time point-specific cold-responsive genes, while those in other cells indicate the number of responsive genes shared between two time points. For example, the number eight in the top left cell indicates that there are eight genes that are up-regulated at both 30 min and 24 hrs. (**B, C)** Biological process GO terms that are significantly enriched (q ≤ 0.05) for genes that are down-regulated (**B**) or up-regulated (**C**) at different time points. Color scale: - log_10_(q) for over-representative GO terms, and log_10_(q) for under-representative GO terms.

**S2 figure:** Feature selection. Graphs show the relationship between the F1score and the number of features in time point models distinguishing genes up-regulated (left panel) or down-regulated (right panel) after cold treatment from non-responsive genes. The trends were fitted using the Michaelis-Menten Equation.

**S3 figure:** pCREs that were identified by models that predicted genes up-regulated at 2~5 time points and their resemblances with known cold-CREs.

**S4 figure:** Heatmap showing feature importance in the cold-TFBM models. Color scale and number in the cell represents the importance rank of features that had positive Gini indexes, the darker color and smaller number, the more important a feature was. Gray color indicates that the Gini index for the feature was negative.

**S5 figure:** A dendrogram showing relationships among general and time point-specific pCREs based on sequence similarities. The dendrogram is clustered based on the similarity threshold of 0.39.

**S6 figure:** Heatmaps showing the differential expression of genes that contain pCREs from different pCRE clusters at different time points after cold treatment. Color scale indicates log fold change values.

**S7 figure:** Frequency of expression profiles (e.g., NNUDNN, y-axis) that are shown by genes that contain pCREs of different pCRE clusters (x-axis). Color scale indicates log_2_(counts) of genes showing the expression profile. U: up-regulated; D: down-regulated; N: non-responsive.

**S8 figure:** Heatmap showing enriched expression profiles (e.g., NNUDNN, y-axis) for genes that contain pCREs of a particular pCRE cluster (x-axis). The color scale represents the log odds ratio, which was calculated as ratios between positive and negative cases.

## SUPPLEMENTAL TABLES

**S1 table:** Metadata of the transcriptome sequences used in this study.

**S2 table:** Number of features selected in the feature selection processes and the best threshold used in different time point models.

**S3 table:** Enrichment *p*-values, feature importance scores, feature importance ranks, and summary of the similarity between pCREs and in-vitro transcription factor binding site data of pCREs in different time point models.

**S4 table:** Information on Transcription Factor Binding Sites (TFBS) of the transcription factors that are known to regulate cold stress response. The table only includes the TFBS whose position weight matrix information was available.

* TFs whose binding sites were similar to the pCRE recovered from our time point models.

^**Y**^ Binding site information was not available in DAP-seq or CISBP datasets

## AUTHOR CONTRIBUTIONS

TR, PW, and SS conceptualized and designed the study. TR and BB acquired and analyzed the data. TR, BB, and PW wrote the original draft of the manuscript. SS and PW supervised the study. All authors read, revised, and approved the final manuscript.

## ACKNOWLEDGEMENT

We thank Kenia Segura Abá, Melissa Lehti-Shiu, Serena Lotreck, and Ally Schumacher for their feedback on the figures and discussion. This work was mainly supported by the US Department of Energy Great Lakes Bioenergy Research Center (BER DE-SC0018409) and in part by the National Science Foundation (DGE-1828149; IOS-2107215; MCB-2210431) to SHS.

