## Supplemental table 4 for "Temporal Regulation of Cold Transcriptional Response in Switchgrass"

**Table S3:** Information on Transcription Factor Binding Sites (TFBS) of the transcription factors that are known to regulate cold stress response. The table only includes the TFBS whose position weight matrix information was available.

| **Transcription Factor** | **Binding Site** | **Cold Response Pathway** | **Citation** |
| --- | --- | --- | --- |
| ABR1* | APETALA2 (AP2) domain (AP2/ERF domain) | CBL9-CIPK3-ABR1 pathway  (Abscisic acid (ABA) signaling pathway) | 29, 34, 40, 41 |
| ANAC062 (NAC062/ANAC062/NTL6) | TT(ACG)CTT | Unfolded protein response pathway (ER) ABA regulation pathway | 27, 43, 58 |
| ATHB2 (HAT4) | CAAT(G/C)ATTG | Auxin Response Pathway | 44, 46 |
| AXR5 (IAA1) **^Ψ^** | NA | Auxin Response Pathway | 56 |
| AZF2 | A(G/C)T repeats (promoter regions of SAUR63 and SAUR20) | Auxin Response Pathway | 22 |
| BZR1* | CGTG(T/C)G | BR signal transduction pathway  (Brassinosteroid signalling) | 13, 52 |
| CAMTA1* | CGCG-box (vCGCGb), CM2 | CBF-COR pathway Salicylic Acid (SA) Immunity Pathway | 7, 19, 20 |
| CAMTA5* | CM2 (5'-CCGCGT-3') | CBF-COR pathway | 7, 20 |
| CBF1* | CRT/DRE (CRT (C-repeat)/DRE (dehydration-responsive element)) - rCCGAC | CBF-COR signaling pathway | 6, 10 |
| CBF2 | rCCGAC | CBF-COR signaling pathway | 35 |
| CBF3* | CRT/DRE | CBF-COR signaling pathway | 6, 10 |
| CIR1 (RVE2)**^Ψ^** | EE (Evening Element) | Circadian Regulation | 11, 59 |
| CRF2 | PIN CYTOKININ RESPONSE ELEMENT (PCRE) GCC-box | Root Development Cytokinin Signaling Pathway Auxin Transport Pathway | 17, 31, 38, 39, 45 |
| CRF3 | PIN CYTOKININ RESPONSE ELEMENT (PCRE) GCC-box | Root Development Cytokinin Signaling Pathway Auxin Transport Pathway | 17, 31, 38, 39, 45 |
| CZF1 (ZFAR1) **^Ψ^** | NA | CBF-COR Pathway | 25 |
| CZF2 | M00861_2.00 | CBF-COR Pathway | 25 |
| DEAR1**^Ψ^** | NA | CBF-COR Pathway Ethylene/Jasmonic Acid (JA) Pathway | 25, 26 |
| DOF1.10**^Ψ^** | AAAG/CTTT | NA | 12, 55 |
| DOF1.3**^Ψ^** | AAAG/CTTT | NA | 12, 55 |
| DREB30 | C-repeat CRT/dehydration responsive element (DRE) motif | CBF-COR Pathway ABA-Independent Signal Transduction Pathway | 1, 25 |
| DREB26* | C- repeat or dehydration response element (DRE) | Stress Tolerance Pathway (Arabidopsis) Early Dehydration Signaling ABA-Independent Signal Transduction Pathway | 14, 16, 23, 47 |
| ERF11 | GCC box (AGCCGCC) C- repeat or dehydration response element (DRE) | Gibberellic Acid (GA) Response Pathway Ethylene Signaling Pathway | 1, 24, 60 |
| ERF115* | GCC box (AGCCGCC) C- repeat or dehydration response element (DRE) | Ethylene Signal Transduction Pathway | 1, 30, 32 |
| ERF5* | GCC box (AGCCGCC) C- repeat or dehydration response element (DRE) | Ethylene signal transduction pathway | 1, 30, 32 |
| ERF6 | GCC box (AGCCGCC) C- repeat or dehydration response element (DRE) | Leaf Growth  Stress Tolerance  Ethylene Signaling Pathway | 1, 8, 53 |
| HSFC1 | M00868_2.00  M00869_2.00  M06936_2.00  M06937_2.00  M06938_2.00 | CBF-COR Pathway | 9, 25 |
| MYB44 | MYB binding site | MKK4 – MPK3 Signaling Pathway ABA-Mediated Signaling Pathway  (related to responding to abiotic stress and stomatal closure) | 15, 25, 37 |
| MYB7 | MYB binding site | ABA Signaling Pathway | 18, 25 |
| MYB73 | MYB recognition site MYB-binding site core motif ACCTAC | Auxin Signaling Pathway  UV-B Photomorphogenesis ABA Signaling Pathway Salt Stress | 25, 51, 57, 58 |
| RAV1 (EDF1) | C(A/C/G)ACA(N)2–8(C/A/T)ACCTG | Ethylene Response Pathway | 2, 28 |
| WRKY22 | W-box | Salicylic Acid (SA) Signaling Pathway  Jasmonic Acid (JA) Signaling Pathway | 21, 48 |
| WRKY24* | W-box W-box (C/T)TGAC(C/T) or W-box like (TGAC[C/T]) | ABA Signaling Pathway | 48, 49, 54 |
| WRKY30* | W-box W-box (C/T)TGAC(C/T) or W-box like (TGAC[C/T]) | JA Signaling Pathway SA Signaling Pathway | 36, 48, 49 |
| WRKY33 | W-box | MKK4/MKK5/MKK9-MPK3/MPK6 Pathway SA-Related Host Response Pathway  JA Response Pathway | 3, 5, 48 |
| WRKY40 | W-box | JA Signaling Pathway SA Signaling Pathway | 33, 42, 48 |
| ZAT10 (STZ) | M00861_2.00, M01168_2.00,  M01852_2.00,  M01854_2.00,  M01860_2.00, | CBF-COR Pathway | 25 |
| ZAT12 | M01859_2.00 | CBF-COR Pathway iron (Fe) Deficiency Response | 4, 25, 50 |
| ZF | M01167_2.00 | CBF-COR pathway iron (Fe) Deficiency Response | 25 |

* The TF found in our pCRE list

**^Ψ^** Binding site information was not available in DAP seq or CISBP datasets

^1^ Agarwal, P.K., Agarwal, P., Reddy, M.K. & Sopory, S.K., 2006. Role of DREB transcription factors in abiotic and biotic stress tolerance in plants. *Plant Cell Reports*, 25, pp.1263–1274.

^2^ Alonso J.M., Stepanova, A.N., Leisse, T.J., Kim, C.J., Chen, H., Shinn, P., et al., 2003. Genome-wide insertional mutagenesis of *Arabidopsis thaliana*. *Science*, 301(5633), pp.653–657.

^3^ Birkenbihl, R.P., Diezel, C. & Somssich, I.E., 2012. Arabidopsis WRKY33 is a key transcriptional regulator of hormonal and metabolic responses toward *botrytis cinerea* infection . *Plant Physiology*, 159(1), pp.266–285.

^4^ Brumbarova, T., Le, C.T.T., Ivanov, R. & Bauer, P., 2016. Regulation of ZAT12 protein stability: The role of hydrogen peroxide. *Plant Signaling & Behavior*, 11(2).

^5^ Datta, R., Kumar, D., Sultana, A., Hazra, S., Bhattacharyya, D. & Chattopadhyay, S., 2015. Glutathione regulates 1-Aminocyclopropane-1-Carboxylate synthase transcription via WRKY33 and 1-Aminocyclopropane-1-Carboxylate Oxidase by modulating messenger RNA stability to induce ethylene synthesis during stress. *Plant Physiology*, 169(4), pp.2963–2981.

^6^ Ding, Y., Shi, Y. & Yang, S., 2019. Advances and challenges in uncovering cold tolerance regulatory mechanisms in plants. *New Phytologist*, 222(4), pp.1690–1704.

^7^ Doherty, C.J., Van Buskirk, H.A., Myers, S.J. & Thomashow, M.F., 2009. Roles for *Arabidopsis* CAMTA transcription factors in cold-regulated gene expression and freezing tolerance. *The Plant Cell*, 21(3), pp.972–984.

^8^ Dubois, M., Van den Broeck, L., Claeys, H., Van Vlierberghe, K., Matsui, M. & Inzé, D., 2015. The ETHYLENE RESPONSE FACTORs ERF6 and ERF11 antagonistically regulate mannitol-induced growth inhibition in *Arabidopsis. Plant Physiology, 169(1), pp.166–179.*

^9^ Franco-Zorrilla, J.M., López-Vidriero, I., Carrasco, J.L., Godoy, M., Vera, P. & Solano, R., 2014. DNA-binding specificities of plant transcription factors and their potential to define target genes. *Proceedings of the National Academy of Sciences*, 111(6), pp.2367–2372.

^10^ Gilmour, S.J., Zarka, D.G., Stockinger, E.J., Salazar, M.P., Houghton, J.M. & Thomashow, M.F., 1998. Low temperature regulation of the Arabidopsis CBF family of AP2 transcriptional activators as an early step in cold-induced COR gene expression. *The Plant Journal*, 16(4), pp.433–442.

^11^ Gong, W., He, K., Covington, M., Dinesh-Kumar, S.P., Snyder, M., Harmer, S.L., et al., 2008. The development of protein microarrays and their applications in DNA–protein and protein–protein interaction analyses of Arabidopsis transcription factors. *Molecular Plant*, 1(1), pp.27–41.

^12^ Gupta, S., Malviya, N., Kushwaha, H., Nasim, J., Bisht, N.C., Singh, V.K. et al., 2015. Insights into structural and functional diversity of Dof (DNA binding with one finger) transcription factor. *Planta*, 241, pp.549–562.

^13^ He, J.-X., Gendron, J.M., Sun, Y., Gampala, S.S.L., Gendron, N., Qing Sun, C. et al., 2005. BZR1 is a transcriptional repressor with dual roles in brassinosteroid homeostasis and growth responses. *Science*, 307(5715), pp.1634–1638.

^14^ Huang, X., Song, X., Chen, R., Zhang, B., Li, C., Liang, Y. et al., 2020. Genome-wide analysis of the DREB subfamily in *Saccharum spontaneum reveals their functional divergence during cold and drought stresses. Frontiers in Genetics, 10.*

^15^ Jung, C., Sio, J.S., Han, S.W., Koo, Y.J., Kim, C.H. & Song, S.I. et al., 2007. Overexpression of *AtMYB44* enhances stomatal closure to confer abiotic stress tolerance in transgenic *Arabidopsis*. *Plant Physiology*, 146(2), pp.323–324.

^16^ Kazama, D., Itakura, M., Kurusu, T., Mitsuda, N., Ohme-Takagi, M. & Tada, Y., 2013. Identification of chimeric repressors that confer salt and osmotic stress tolerance in Arabidopsis. *Plants*, 2(4), pp.769–785.

^17^ Kim, J., 2016. CYTOKININ RESPONSE FACTORs gating environmental signals and hormones. *Trends in Plant Science*, 21(12), pp.993–996.

^18^ Kim, J.H., Hyun, W.Y., Nguyen, H.N., Jeong, C.Y., Xiong, L., Hong, S.-W. et al., 2014. AtMyb7, a subgroup 4 R2R3 Myb, negatively regulates ABA-induced inhibition of seed germination by blocking the expression of the bZIP transcription factor ABI5. *Plant, Cell & Environment*, 38(3), pp.559–571.

^19^ Kim, Y.S., Park, S., Gilmour, S.J. & Thomashow, M.F. 2013. Roles of CAMTA transcription factors and salicylic acid in configuring the low-temperature transcriptome and freezing tolerance of Arabidopsis. *The Plant Journal*, 75(3), pp.364–376.

^20^ Kim, Y.S., An, C., Park, S., Gilmour, S.J., Wang, L., Renna, L. et al., 2017. CAMTA-mediated regulation of salicylic acid immunity pathway genes in arabidopsis exposed to low temperature and pathogen infection. *The Plant Cell*, 29(10), pp.2465–2477.

^21^ Kloth, K.J., Wiegers, G.L., Busscher-Lange, J., van Haarst, J.C., Kruijer, W., Bouwmeester, H.J., et al., 2016. AtWRKY22 promotes susceptibility to aphids and modulates salicylic acid and jasmonic acid signalling. *Journal of Experimental Botany*, 67(11), pp.3383–3396.

^22^ Kodaira, K.-S., Qin, F., Phan Tran, L.-S., Maruyama, K., Kidokoro, S., Fujita, Y. et al., 2011. Arabidopsis Cys2/his2 zinc-finger proteins AZF1 and AZF2 negatively regulate abscisic acid-repressive and auxin-inducible genes under abiotic stress conditions  . *Plant Physiology*, 157(2), pp.742–756.

^23^ Krishnaswamy, S., Verma, S., Rahman, M.H. & Kav, N.N.V., 2010. Functional characterization of four APETALA2-family genes (RAP2.6, RAP2.6L, DREB19 and DREB26) in Arabidopsis. *Plant Molecular Biology*, 75, pp.107–127.

^24^ Li, Z., Zhang, L., Yu, Y., Quan, R., Zhang, Z., Zhang, H. et al., 2011. The ethylene response factor AtERF11 that is transcriptionally modulated by the bZIP transcription factor HY5 is a crucial repressor for ethylene biosynthesis in Arabidopsis. *The Plant Journal*, 68(1), pp.88–99.

^25^ Liu, Y., Dang, P., Liu, L. & He, C., 2019. Cold acclimation by the CBF–COR pathway in a changing climate: Lessons from *Arabidopsis thaliana*. *Plant Cell Reports*, 38(5), pp.511–519.

^26^ Maruyama, Y., Yamoto, N., Suzuki, Y., Chiba, Y., Yamazaki, K.-I., Sato, T., et al., 2013. The Arabidopsis transcriptional repressor ERF9 participates in resistance against necrotrophic fungi. *Plant Science*, 213, pp.79–87.

^27^ Mathew, I.E. & Agarwal, P., 2018. May the fittest protein evolve: Favoring the plant-specific origin and expansion of NAC transcription factors. *BioEssays*, 40(8).

^28^ Matías-Hernández, L., Aguilar-Jaramillo, A.E., Marín-González, E., Suárez-López, P. & Pelaz, S., 2014. RAV Genes: Regulation of floral induction and beyond. *Annals of Botany*, 114(7), pp.1459–1470.

^29^ Mishra, M., Kanwar, P., Singh, A., Pandey, A., Kapoor, S. & Pandey, G.K., 2013. Plant omics: Genome-wide analysis of Aba Repressor1 (*abr1*) related genes in rice during abiotic stress and development. *OMICS: A Journal of Integrative Biology*, 17(8), pp.439–450.

^30^ Moffat, C.S., Ingle, R.A., Wathugala, D.L., Saunders, N.J., Knight, H. & Knight, M.R., 2012. ERF5 and ERF6 play redundant roles as positive regulators of JA/Et-mediated defense against *Botrytis cinerea in Arabidopsis. PLoS ONE, 7(4).*

^31^ Oh, E., Kang, H., Yamaguchi, S., Park, J., Lee, D., Kamiya, Y. et al., 2009. Genome-wide analysis of genes targeted by PHYTOCHROME INTERACTING FACTOR 3-LIKE5 during seed germination in *Arabidopsis*. *The Plant Cell*, 21(2), pp.403–419.

^32^ Ohme-Takagi, M. & Shinshi, H., 1995. Ethylene-inducible DNA binding proteins that interact with an ethylene-responsive element. *The Plant Cell*, 7(2), pp.173–182.

^33^ Pandey, G.K., Grant, J.J., Cheong, Y.H., Kim, B.G., Li, L., & Luan, S., 2005. ABR1, an APETALA2-domain transcription factor that functions as a repressor of ABA response in Arabidopsis. *Plant Physiology*, 139(3), pp.1185–1193.

^34^ Pandey, S.P., Roccaro, M., Schön, M., Logemann, E. & Somssich, I.E., 2010. Transcriptional reprogramming regulated by WRKY18 and WRKY40 facilitates powdery mildew infection of Arabidopsis. *The Plant Journal*, 64(6), pp.912–923.

^35^ Park, S., Gilmour, S.J., Grumet, R. & Thomashow, M.F., 2018. CBF-dependent and CBF-independent regulatory pathways contribute to the differences in freezing tolerance and cold-regulated gene expression of two Arabidopsis ecotypes locally adapted to sites in Sweden and Italy. *PLOS ONE*, 13(12).

^36^ Peng, X., Hu, Y., Tang, X., Zhou, P., Deng, X., Wang, H., et al., 2012. Constitutive expression of rice WRKY30 gene increases the endogenous jasmonic acid accumulation, PR gene expression and resistance to fungal pathogens in Rice. *Planta*, 236(5), pp.1485–1498.

^37^ Persak, H. & Pitzschke, A., 2013. Tight interconnection and multi-level control of Arabidopsis MYB44 in MAPK cascade signalling. *PLoS ONE*, 8(2).

^38^ Rashotte, A.M. & Goertzen, L.R., 2010. The CRF domain defines cytokinin response factor proteins in plants. *BMC Plant Biology*, 10(1), p.74.

^39^ Rashotte, A.M., Mason, A.M., Hutchison, C.E., Ferreira, F.J., Schaller, G.E. & Kieber, J.J., 2006. A subset of *Arabidopsis* AP2 transcription factors mediates cytokinin responses in concert with a two-component pathway. *Proceedings of the National Academy of Sciences*, 103(29), pp.11081–11085.

^40^ Sanyal, S.K., Kanwar, P., Yadav, A.K., Sharma, C., Kumar, A., & Pandey, G.K., 2017. Arabidopsis CBL interacting protein kinase 3 interacts with ABR1, an APETALA2 domain transcription factor, to regulate ABA responses. *Plant Science*, 254, pp.48–59.

^41^ Schreiber, K.J., Hassan, J.A. & Lewis, J.D., 2021. Arabidopsis abscisic acid repressor 1 is a susceptibility hub that interacts with multiple *pseudomonas syringae* effectors. *The Plant Journal*, 105(5), pp.1274–1292.

^42^ Schön, M., Töller, A., Diezel, C., Roth, C., Westphal, L., Wiermer, M., et al., 2013. Analyses of *WRKY18 wrky40* plants reveal critical roles of SA/EDS1 signaling and indole-glucosinolate biosynthesis for *golovinomyces orontii* resistance and a loss-of resistance towards *pseudomonas syringae* pv. *tomato* avrrps4. *Molecular Plant-Microbe Interactions®*, 26(7), pp.758–767.

^43^ Seo, P.J. & Park, C.-M., 2010. A membrane-bound NAC transcription factor as an integrator of biotic and abiotic stress signals. *Plant Signaling & Behavior*, 5(5), pp.481–483.

^44^ Sessa, G., Morelli, G. & Ruberti, I., 1993. The ATHB-1 and −2 HD-zip domains homodimerize forming complexes of different DNA binding specificities. *The EMBO Journal*, 12(9), pp.3507–3517.

^45^ Šimášková, M., O’Brien, J.A., Khan, M., Van Noorden, G., Ötvös, K., Vieten, A., et al., 2015. Cytokinin response factors regulate pin-formed auxin transporters. *Nature Communications*, 6(1).

^46^ Steindler, C., Matteucci, A., Sessa, G., Weimar, T., Ohgishi, M., Aoyama, T., et al., 1999. Shade avoidance responses are mediated by the ATHB-2 HD-zip protein, a negative regulator of gene expression. *Development*, 126(19), pp.4235–4245.

^47^ Urano, K., Maruyama, K., Jikumaru, Y., Kamiya, Y., Yamaguchi-Shinozaki, K. & Shinozaki, K., 2016. Analysis of plant hormone profiles in response to moderate dehydration stress. *The Plant Journal*, 90(1), pp.17–36.

^48^ Viana, V.E., Busanello, C., Carlos da Maia, L., Pegoraro, C., Costa de Oliveira, A, 2018. Activation of rice WRKY transcription factors: An army of stress fighting soldiers? *Current Opinion in Plant Biology*, 45(Part B), pp.268–275.

^49^ Viana, V.E., Carlos da Maia, L., Busanello, C., Pegoraro, C. & Costa de Oliveira, A., 2021. When Rice gets the Chills: Comparative transcriptome profiling at germination shows WRKY transcription factor responses. *Plant Biology*, 23(S1), pp.100–112.

^50^ Vogel, J.T., Zarka, D.G., Van Buskirk, H.A., Fowler, S.G., Thomashow, M.F., 2004. Roles of the CBF2 and ZAT12 transcription factors in configuring the low temperature transcriptome of Arabidopsis. *The Plant Journal*, 41(2), pp.195–211.

^51^ Wang, L., Qiu, T., Yue, J., Guo, N., He, Y., Han, X. et al., 2021. *Arabidopsis ADF1* is regulated by MYB73 and is involved in response to salt stress affecting actin filament organization. *Plant and Cell Physiology*, 62(9), pp.1387–1395.

^52^ Wang, Z.-Y., Nakano, T., Gendron, J., He, J., Chen, M., Vafeados, D. et al., 2002. Nuclear-localized BZR1 mediates brassinosteroid-induced growth and feedback suppression of brassinosteroid biosynthesis. *Developmental Cell*, 2(4), pp.505–513.

^53^ Warmerdam, S., Sterken, M.G., Van Schaik, C., Oortwijn, M.E.P., Lozano-Torres, J.L., Bakker, J. et al., 2018. Mediator of tolerance to abiotic stress ERF6 regulates susceptibility of *Arabidopsis to Meloidogyne incognita. Molecular Plant Pathology, 20(1), pp.137–152.*

^54^ Xie, Z., Zhang, Z.-L., Zou, X., Huang, J., Ruas, P., Thompson, D., et al., 2005. Annotations and functional analyses of the Rice *WRKY* gene superfamily reveal positive and negative regulators of abscisic acid signaling in Aleurone cells . *Plant Physiology*, 137(1), pp.176–189.

^55^ Yanagisawa, S., 2002. The Dof family of plant transcription factors. *Trends in Plant Science*, 7(12), pp.555–560.

^56^ Yang, X., Lee, S., So, J.-H., Dharmasiri, S., Dharmasiri, N., Ge, L., Jensen, C., et al., 2004. The IAA1 protein is encoded by AXR5 and is a substrate of SCFTIR1. *The Plant Journal*, 40(5), pp.772–782.

^57^ Yang, Y., Zhang, L., Chen, P., Liang, T., Li, X. & Liu, H., 2019. UV‐B photoreceptor UVR8 interacts with MYB73/MYB77 to regulate auxin responses and lateral root development. *The EMBO Journal*, 39(2).

^58^ Yang, Z.-T., Lu, S.-J., Wang, M.-J., Bi, D.-L., Sun, L., Zhou, S.-F., et al., 2014. A plasma membrane-tethered transcription factor, NAC062/ANAC062/NTL6, mediates the unfolded protein response in Arabidopsis. *The Plant Journal*, 79(6), pp.1033–1043.

^59^ Zhang, X., Chen, Y., Wang, Z.-Y., Chen, Z., Gu, H., Qu, L.-J., 2007. Constitutive expression of CIR1 (RVE2) affects several circadian-regulated processes and seed germination in Arabidopsis. *The Plant Journal*, 51(3), pp.512–525.

^60^ Zhao, Y. et al., 2014. The ABA receptor PYL8 promotes lateral root growth by enhancing MYB77-dependent transcription of Auxin-responsive genes. *Science Signaling*, 7(328).

^61^ Zhou, X., Zhang, Z.-L., Park, J., Tyler, L., Yusuke, J., Qiu, K. et al., 2016. The ERF11 transcription factor promotes internode elongation by activating gibberellin biosynthesis and signaling. *Plant Physiology*, 171(4), pp.2760–2770.
